# Sequence assignment validation in cryo-EM models with checkMySequence

**DOI:** 10.1101/2022.01.04.474974

**Authors:** Grzegorz Chojnowski

## Abstract

The availability of new AI-based protein structure prediction tools radically changed the way cryo-EM maps are interpreted, but it has not eliminated the challenges of map interpretation faced by a microscopist. Models will continue to be locally rebuilt and refined using interactive tools. This inevitably results in occasional errors, among which register-shifts remain one of the most difficult to identify and correct. Here we introduce *checkMySequence*; a fast, fully automated and parameter-free method for detecting register-shifts in protein models built into cryo-EM maps. We show that the method can assist model building in cases where poorer map resolution hinders visual interpretation. We also show that *checkMySequence* could have helped avoid a widely discussed sequence register error in a model of SARS-CoV-2 RNA-dependent RNA polymerase that was originally detected thanks to a visual residue-by-residue inspection by members of the structural biology community.

**Synopsis:** We present a new method, *checkMySequence*, for fast and automated detection of register errors in protein models built into cryo-EM reconstructions.

## 1. Introduction

Five years after the resolution revolution in cryogenic electron-microscopy (cryo-EM) began, we are witnessing another revolution; in the accuracy of protein structure prediction techniques. The former paved the way to the structure determination of large macromolecular complexes, in the absence of crystals and at a level of detail enabling studying of biological processes at atomic scale (Kühlbrandt, 2014; Kokic *et al*., 2021). The latter provided means for accurate and widely accessible structure prediction of protein structures. Although the release of Artificial Intelligence (AI)-based AlphaFold 2 (AF2) (Jumper *et al*., 2021) and the similar RoseTTAFold (Baek *et al*.,2021) represent dramatic progress in the way protein structures are predicted, they did not solve the problem of protein structure determination. Structural studies of multimeric, highly dynamic complexes or specific ligand-bound states of proteins still require experimental approaches and from this perspective the new structure prediction approaches perfectly complement cryo-EM (Perrakis & Sixma, 2021). Nevertheless, the new predictive methods have dramatically changed the way cryo-EM reconstructions are interpreted (Mosalaganti *et al*., 2021).

With the availability of accurate protein structure prediction tools, interpretation of most cryo-EM reconstructions doesn’t require *de novo* model tracing (e.g. with ARP/wARP (Chojnowski, Sobolev *et al*., 2021), buccaneer (Hoh *et al*., 2020), phenix.map_to_model (Terwilliger *et al*.,2018)) or parallel structure determination of model components using X-ray crystallography (Beckham *et al*., 2021). Instead, one can follow an approach often used in cryo-EM at lower resolutions, where whole models are assembled from experimentally or theoretically determined structures of their components (Allegretti *et al*., 2020). The structure assembly procedure, however, does not eliminate the need for interactive (or “manual”) model rebuilding. This is still required in regions where a reliable structure cannot be predicted due to low sequence coverage, or was predicted in a conformational state incompatible with a target (Perrakis & Sixma, 2021). Similarly, interfaces within homo- or hetero-multimers often cannot be reliably modelled, even with an excellent, community driven AF2-extension ColabFold (Mirdita *et al*., 2021), or recently released AlphaFold-Multimer (Evans *et al*., 2021). In such cases tools like COOT (Casañal *et al*.,2020) or ISOLDE (Croll, 2018) make the interactive model refinement and rebuilding relatively fast and simple, but still requiring subjective visual interpretation of a map by a user. This process, although significantly simplified due to the availability of reliably predicted initial models, inevitably results in sporadic errors.

Errors may occur in every structure, regardless of the resolution, or experimentalist’s best efforts. However, they seem to be more common in cryo-EM where models are often built quickly, under pressure, and into reconstructions spanning a wide range of local resolutions. Although many issues can be corrected automatically (Joosten *et al*., 2014; Liebschner *et al*., 2021), visual residue-by-residue inspection by an experienced structural biologist remains the best way to judge the quality of a model. This, however, is time consuming and requires a level of expertise that is rarely available (Croll, Williams *et al*., 2021). Therefore, computational expert-systems for model-validation like Molprobity (Chen *et al*., 2010) are indispensable in routine detection of modelling errors. Even these tools, however, usually require experience in separating severe problems that must be corrected from those unusual features that may be left in a model. Moreover, finding an optimal way of correcting an issue is not always straightforward.

In cryo-EM, one of the most difficult problems to identify and correct are register-shift errors, where residues are systematically assigned an identity of a residue a few amino acids up or down in sequence. When resolution allows, register-shifts can be identified using aromatic residues, which are usually well-resolved in the density. This can be conveniently done using a dedicated, interactive tool implemented in ISOLDE. The process, however, cannot be easily automated as map-fit measures are usually more sensitive to atoms outside density, than density left without a model (Croll, Williams *et al*., 2021). Nevertheless, tools like EM-ringer (Barad *et al*., 2015), Q-score (Pintilie *et al*., 2020), or SMOC (Joseph *et al*., 2016) can in principle be used for detecting register-shifts, even though there are no clear criteria that might be used to translate validation score fluctuations to specific problems in a model. Moreover, these density-fit scores are strongly local-resolution dependent, which may hinder recognition of register-shifts from effects of tracing problems or variations of local-resolution.

Register-shift errors may also have an effect on backbone geometry when a number of side-chains are forced into too small density volumes. It has been shown that these can in principle be detected using CaBLAM (Richardson *et al*., 2018). Moreover, the very source of register shift that is often a backbone tracing issue (e.g. deletion or insertion) can be occasionally detected based on backbone geometry problems (Lawson *et al*., 2021).

Neverthess, to the best of our knowledge, there is no tool available that was developed specifically for automated detection of register-shift errors in macromolecular models. Although available model-geometry and density-fit validation tools can in some cases help to detect these, there are no clear rules-of-thumb that would allow for a selection of troubled regions that need to be carefully checked by a user. Furthermore, none of these tools can automatically suggest a possible fix to a plausible register-shift problem.

Here we present *checkMySequence*, a new tool for automated detection of register-shift errors in cryo-EM models. The method is based on *findMySequence*, a protein sequence identification tool for crystallography and cryo-EM (Chojnowski, Simpkin *et al*., 2022). The *checkMySequence* algorithm uses tools implemented in *findMySequence* to assign input-model fragments to a reference sequence. It identifies regions where the new sequence-assignment challenges the sequence-assignment hypothesis in the input model. This approach provides a conceptually simple, fast and intuitive tool for reliable detection of register-shifts in cryo-EM models including very large macromolecular complexes (e.g. complete ribosomes). If an issue is detected, the method suggests to the user a more plausible sequence assignment. We show that *checkMySequence* can reliably identify register-shift errors in models deposited to the Protein Data Bank (PDB)(Berman *et al*., 2000) that were already reported in the literature and a number of new, previously unidentified ones.

## 2. Model validation with a systematic sequence assignment

The method requires on input a cryo-EM map, corresponding atomic model, and sequences of all the model chains. Initially, for each protein chain in the input model the method identifies a reference sequence. It uses a protocol implemented in the *findMySequence* program (Chojnowski, Simpkin *et al*., 2022) based on a Neural-Network residue-type classifier and HMMER (Eddy, 2011) sequence comparison suite (Fig. 1a). Each chain, for which reference sequence can be identified, is divided into continuous overlapping test-fragments that are systematically assigned to the reference sequence. The program identifies the most plausible assignment of a test-fragment to a reference sequence given residue-type probabilities estimated from a map and backbone coordinates. We assume here that the test-fragments are continuous and ignore all modified residues in the model. To account for variations in the accuracy of residue-type probability estimates and different lengths of reference sequences, for each assignment we estimate a p-value, or a probability that it was observed by chance. Additionally, to compensate for lower local-resolutions, the initial fragment length of 20 residues is increased up to a maximum of 60 residues, if the corresponding sequence assignment p-value exceeds a threshold defined in section 3.1 (Fig. 1b). Depending on the result of the initial reference sequence identification and a test-fragment sequence assignment procedure the following alternative outcomes are possible:

- Reference sequence for a chain cannot be identified; corresponding sequence is missing in input, chain is traced in a very low local-resolution region, or mistraced.
- Sequence assignment of a test-fragment is unreliable (p-value above a threshold defined in section 3.1); there is not enough evidence to confirm or reject corresponding input-model sequence.
- Sequence assignment of a test-fragment is confident (p-value is below the threshold) and assigned sequence agrees with input model; corresponding input-model sequence assignment is confirmed.
- Sequence assignment of a test-fragment is confident (p-value is below the threshold), but assigned sequence doesn’t agree with input model; corresponding input-model sequence may be register-shifted

**Figure 1.**
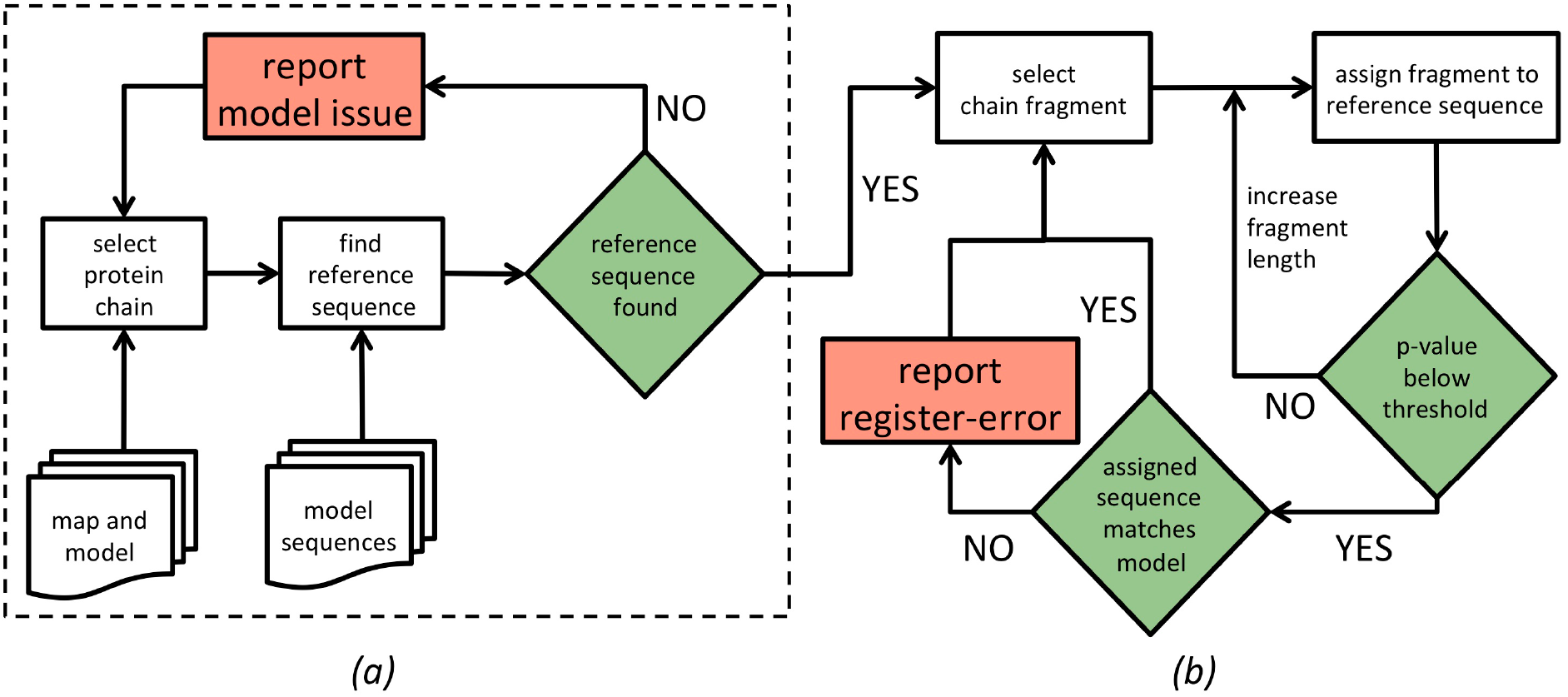
Flowchart of a model-sequence validation procedure. Dashed box (a) encloses the initial steps of reference sequence identification for each protein chain in the input model. If a reference sequence can be identified, the chain is divided into continuous test-fragments that are systematically assigned to the reference sequence (b). The method reports possible issues (red boxes) if a reference sequence cannot be identified, or a sequence assigned to a test-fragment with high confidence (p-value below threshold) doesn’t match the input model.

## 3. Results and discussion

### 3.1. Sequence assignment validation - finding a better hypothesis

Our working hypothesis is that the input model sequence is correct and agrees with an unknown, ground truth reference model. If a result of the sequence assignment procedure is conclusive the hypothesis can be either confirmed or challenged by providing a model that explains cryo-EM map features better. This approach, however, requires clear criteria for assessing statistical significance of the sequence assignment results.

The sequence assignment procedure used in this work has been calibrated to provide a p-value estimate; a probability that the result was obtained by chance. In the current setup, however, we have no means to validate reliability of this in detail as the reference structures available in PDB may and do occasionally contain sequence register errors that can be identified and corrected only by detailed inspection by an experienced modeller. Indeed, in a recent group effort “Coronavirus Structural Task Force’’ members were able to identify multiple tracing and sequence assignment errors in a relatively small, representative set of PDB deposited *Sarbecovirus* protein models (Croll, Diederichs *et al*., 2021).

To address the issue of a benchmark set reliability, we decided to undertake a large-scale, non-parametric approach assuming that errors in deposited PDB models are scarce and will weakly affect overall conclusions. We tested agreement with reference models of sequences assigned to continuous protein chain fragments of 20 amino acids systematically selected from 796 protein models from the benchmark set described in Materials and Methods.

For a total number of 166,713 protein chain test-fragments for which a reference sequence could be identified (see Materials and Methods for details), the assigned sequence matched the corresponding model in 156,091 (94%) and differed in 10,622 (6%) of cases. Protein chain fragments with assigned sequence matching and different from reference are well separated by corresponding p-value (Fig. 2a). Indeed, a one-sided 99.5% confidence interval for fragments with sequence assignment that doesn’t match the reference (dashed line on Fig. 2a) corresponds to 13% of cases with matching sequences. The relatively large number of fragments assigned a correct sequence with high p-values may be due to the presence of model stretches built into local-resolution regions too low for reliable sequence assignment. Indeed, the distribution of correctly assigned sequences is clearly shifted towards better local-resolutions compared to assignments with incorrect sequence (Fig. 2b). These observations clearly show that the p-value is a reliable criterion of sequence assignment procedure validity. Moreover, as expected, if sequence-assignment errors are present in the benchmark set models, they are relatively rare.

**Figure 2.**
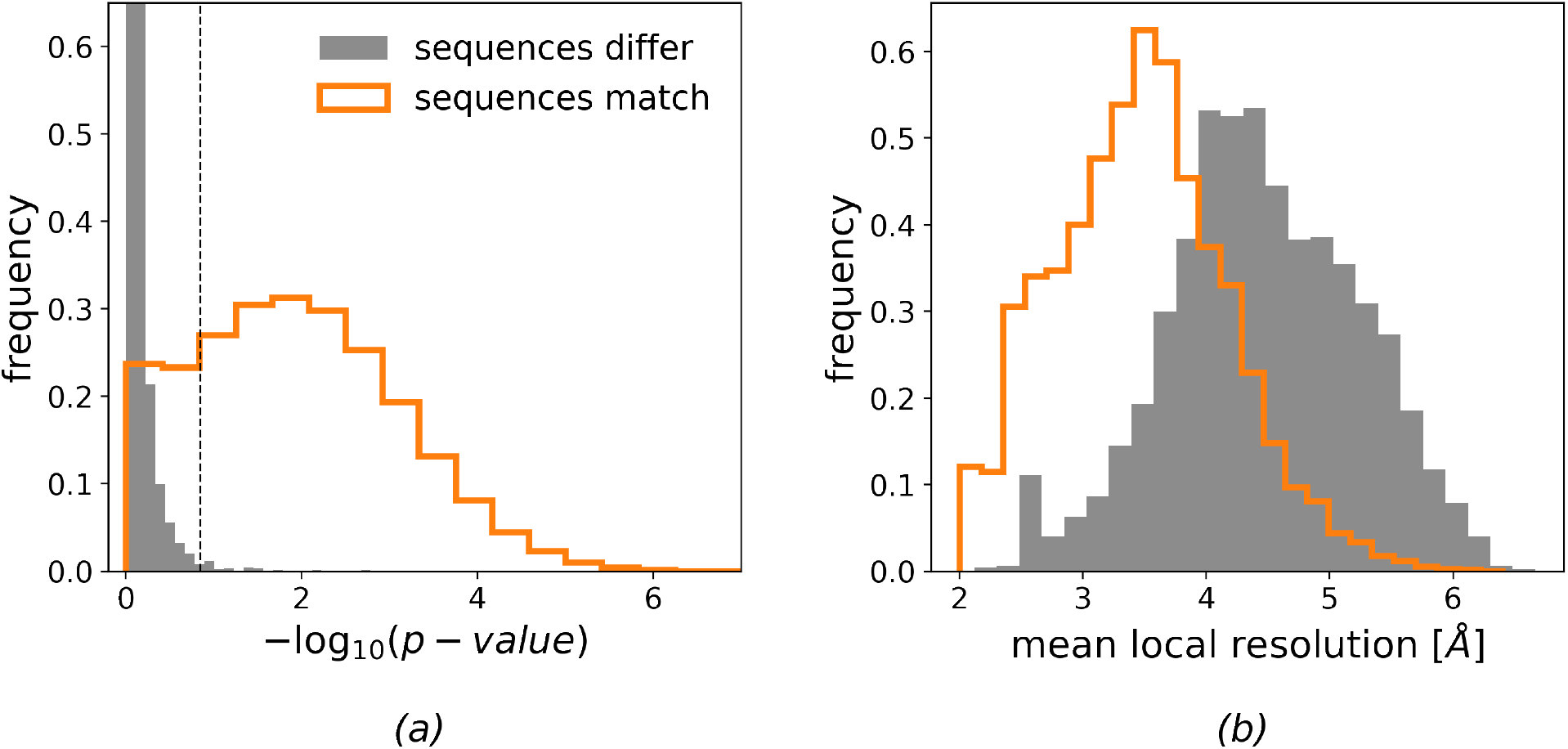
Comparison of sequence assignment results for 166,713 protein chain test-fragments of 20 residues from the benchmark set. Distribution of (a) p-value and (b) mean local resolution of test-fragments, which assigned sequence match and differ from the reference model. Dashed line in panel (a) corresponds to a 99.5% one-sided confidence interval estimated for fragments with an assigned sequence different than reference.

In this work we treat as conclusive sequence assignments with a p-value outside the 99.5% onesided confidence interval estimated for fragments with assigned sequence mismatch (Fig 2a). Although the choice of threshold is arbitrary it corresponds to results that are very rare in our benchmark set and may indicate an outlier. We will show later that indeed many sequence assignments outside this p-value interval are due to plausible reference structure errors.

### 3.2. Compensating low local-resolution

In the previous section we noted that the observed protein fragments with correctly assigned sequence and high p-value may correspond to map regions of local-resolution too low for *de novo* tracing. To further investigate this issue, we plotted sequence assignment p-value as a function of mean local-resolution of corresponding test-fragments. The p-values are clearly lower for test-fragments modelled into lower local-resolution regions (Fig. 3a) and for half of the fragments exceed the threshold defined in the previous section at resolutions as low as 6 Å. At the same time, we observed that p-values for sequence assignment that don’t match the reference are local-resolution independent (Fig. 3b). This clearly indicates that sequence misassignment is often related to intrinsic properties of a model (e.g. tracing errors), not low information content of a corresponding map region. We also observed that the p-value gap between correct and incorrect sequence assignments increases for longer fragments. This can be used to compensate for lower local-resolution.

**Figure 3.**
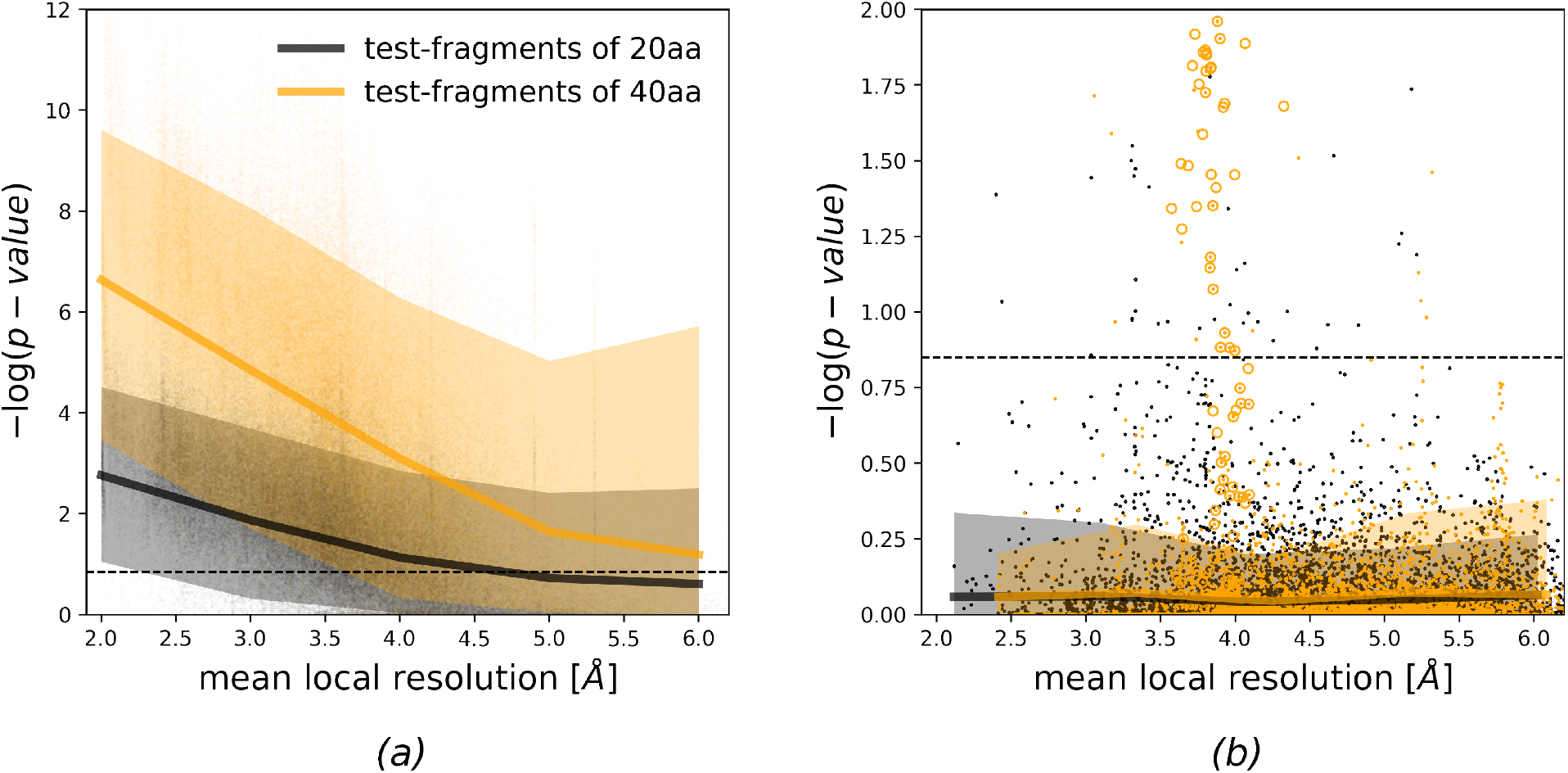
Medians and 90% confidence intervals for sequence assignment p-value as a function of local resolution for protein chain test-fragments of 20 and 40 amino acid residues. Panel (a) shows fragments with sequences matching reference. Fragments for which assigned and reference model sequences differ are presented in panel (b). Dashed line corresponds to a 99.5% one-sided confidence interval estimated for fragments of 20 amino acids with an assigned sequence different from the input-model sequence. Orange circles depict an outlier discussed in the text (cytoplasmic domain of a Transient receptor potential channel, pdb-id 6cv9).

### 3.3. The effect of coordinate errors

To test the validity of our earlier observations in the presence of model errors, we plotted the sequence assignment p-values for protein chain test-fragments at various levels of randomisation of atom coordinates. We observed that the coordinate randomisation increases p-values for all fragments, regardless of whether their assigned sequence matches or doesn’t match the reference model (Fig. 4). As a result, fewer fragments exceed the p-value threshold defined in the previous section (Fig. 2a), which reduces the predictive power of the presented approach. Nevertheless, we observed that the negative effect of coordinate errors can be compensated with longer test-fragments (Fig. 4a).

**Figure 4.**
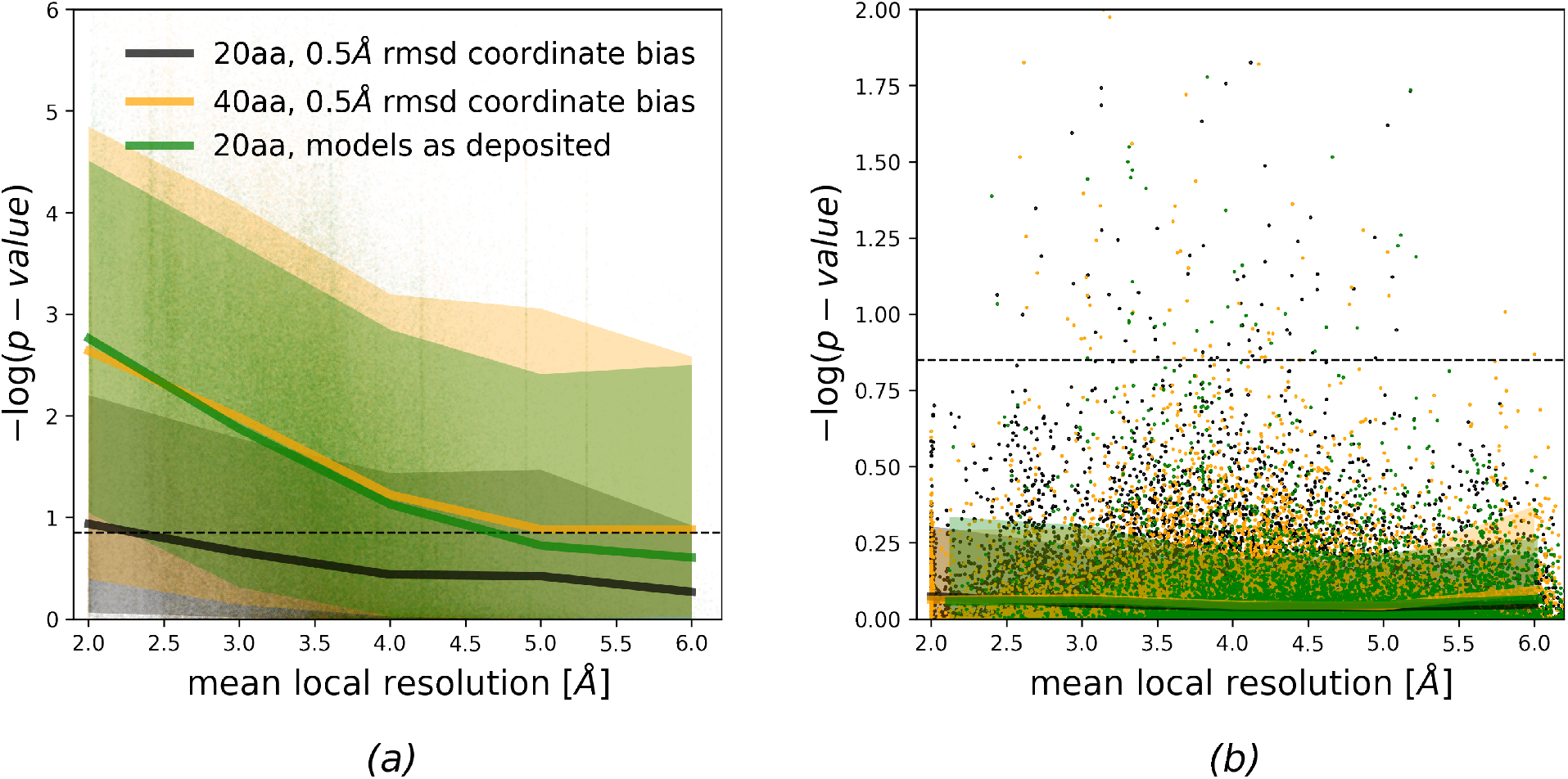
Medians and 90% confidence intervals for sequence assignment p-value as a function of local resolution. Data are shown for protein chain fragments of 20 and 40 aminos acid residues for models as deposited and in the presence of artificial coordinate bias. Panel (a) shows fragments with sequences matching reference. Fragments for which reassigned and reference sequences differ are presented on panel (b). The dashed line corresponds to a 99.5% one-sided confidence interval estimated for fragments of 20 amino acids with an assigned sequence different from the corresponding input-model sequence.

### 3.4. The effect of map sharpening and blurring

In earlier work, we observed that many deposited maps are over-sharpened to a level hindering their interpretation (Chojnowski, Sobolev *et al*., 2021). As maps are often sharpened or blurred by microscopists based on subjective criteria, for example to increase interpretability of specific map regions (Nicholls *et al*., 2018), we decided to investigate the effect of excessive map processing on our sequence assignment procedure.

We observed that the excessive input map sharpening and blurring have a similar influence on sequence assignment p-values (not shown). However, map sharpening (Fig. 5) has more impact on fragments built into high local-resolution regions. We attribute this to the Neural-Network residue-type classifier used in this work, which was trained on an unstratified set of deposited maps. It reflects experimentally observed distribution of local-resolutions, where very high local-resolution regions may be under-represented. We observed that the negative effect of excessive map sharpening or blurring (data for blurred maps was omitted for clarity) can be easily compensated with a longer test-fragment length (Fig. 5a). We did not observe any effect related to map blurring or sharpening for test-fragments with assignment sequence not matching reference model.

**Figure 5.**
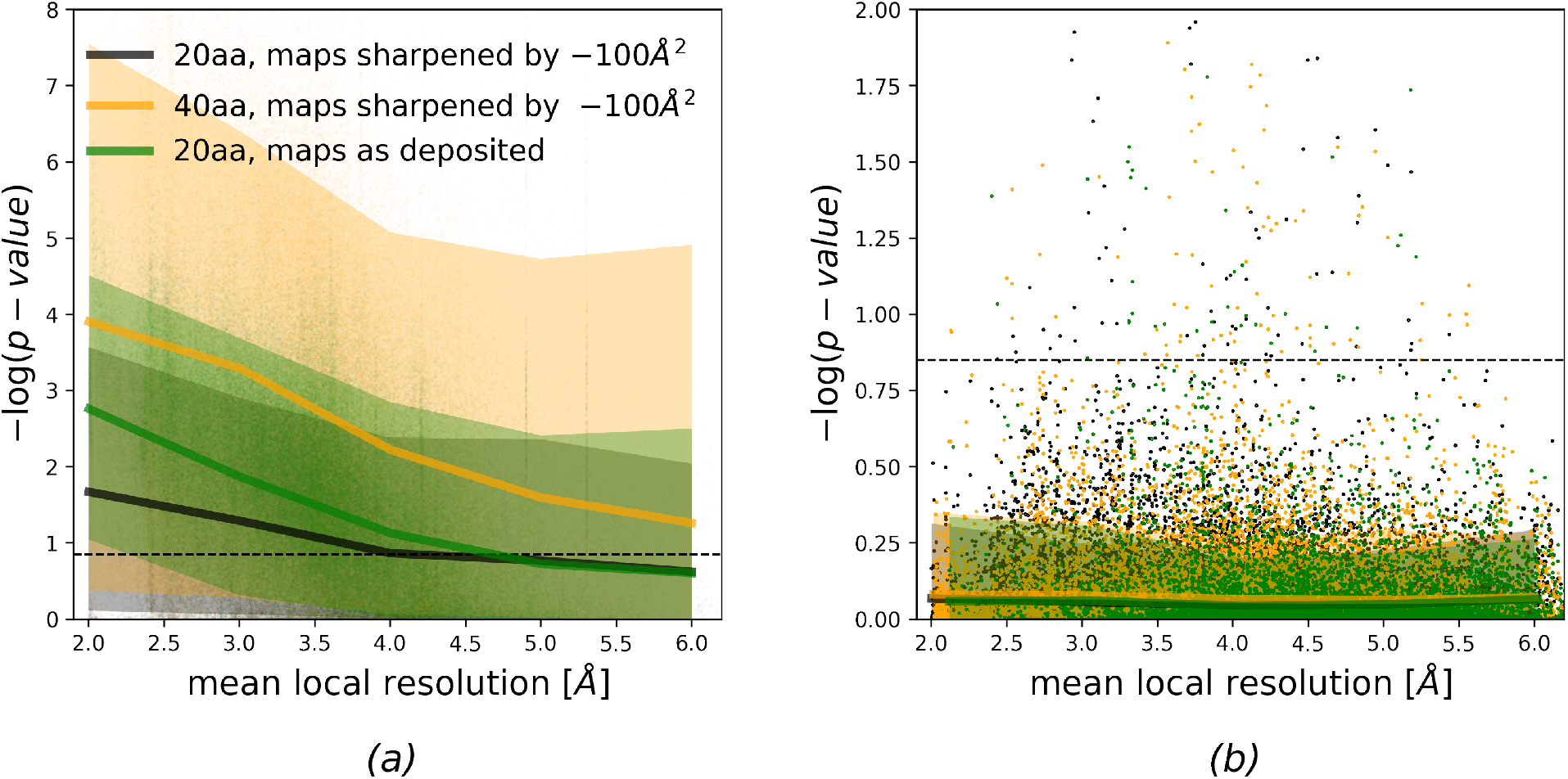
Median and 90% confidence intervals for sequence assignment p-value as a function of local resolution. Data are shown for protein chain fragments of 20 and 40 amino acids and corresponding maps as deposited or sharpened by −100 Å^2^. Panel (a) shows fragments with sequences matching reference. Fragments for which assigned and reference-model sequences differ are presented on panel (b). The dashed line corresponds to a 99.5% one-sided confidence interval estimated for fragments of 20 amino acids with an assigned sequence different from the corresponding input-model sequence.

### 3.5. Reference sequence identification performance

A crucial step in the described method is the identification of reference sequences for each chain in the input model. For this purpose, we use a procedure implemented in *findMySequence*. The program uses a Neural-Network classifier to produce residue-type probability profiles that are further used to query input sequence sets with tools from HMMER suite. Sequence matches identified by HMMER are scored with an E-value, a number of expected hits in a comparable set containing only random sequences. The lower the E-values, the more reliable the corresponding match. Since this approach is inherently very sensitive to map and model quality, we investigated how coordinate errors and excessive map sharpening or blurring affects the performance of the reference sequence identification procedure.

Overall, out of 3378 protein chains longer than 10 residues in our benchmark set, *checkMySequence* correctly identified 3162 (94%). For the remaining 216 structures the program returned no results, which is reported to a user as a plausible error. Indeed, we observed that many of them were clearly shifted outside corresponding cryo-EM reconstructions (for example pdb/emdb-ids: 6vx7/21432, 6vp8/21250, 6tqm/10555, 6xi8/22191).

We observed that the map processing increases HMMER E-values, but the effect of sharpening (negative B-factor correction) is more prominent than blurring (Fig. 6a). There are also a number of structures for which map blurring significantly improves method performance (lower E-values). Moreover, the overall number of 3162 chains in the benchmark set for which reference sequence could have been identified reduces to 3149 for blurred maps and to 2508 for sharpened maps. This is in line with an earlier observation that many deposited maps are heavily over-sharpened (Chojnowski, Sobolev *et al*., 2021). Interestingly, we also observed that the coordinate bias reduces the number of identified sequences less than excessive map sharpening (Fig. 6b). Initial number of recognized chain sequences reduces to 3151, 3100, and 2797 for 0.1 Å, 0.3 Å, and 0.5 Å all-atom rmsd respectively. Nevertheless, the identified sequences were always correct, regardless of map processing or model bias.

**Figure 6.**
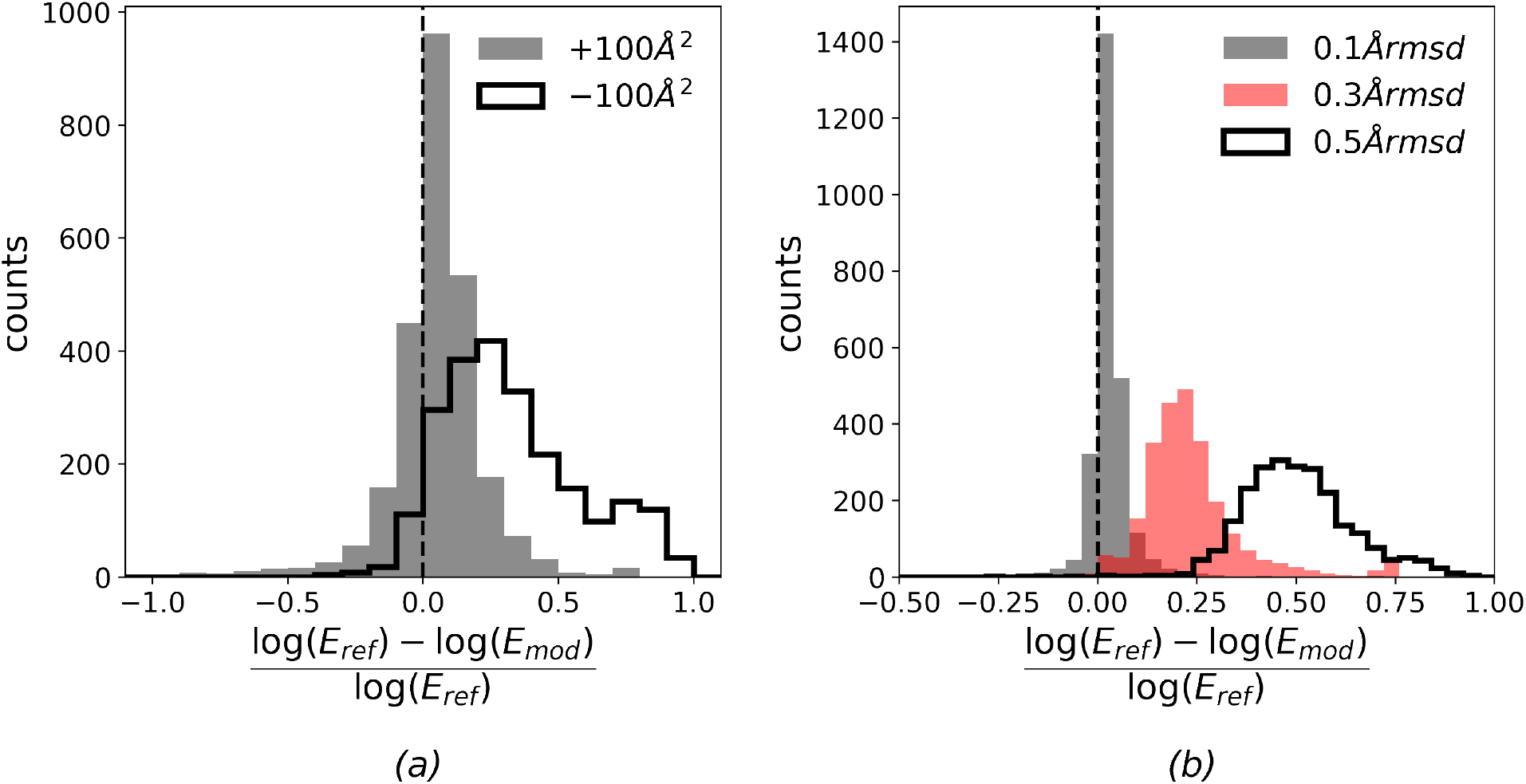
Relative change in E-value estimates for reference sequences identified by *checkMySequence* using HMMER suite. The effect of (a) map blurring or sharpening (B-factor correction +100 and −100 Å^2^ respectively) and (b) model coordinates bias. Eref refers to E-values obtained for reference models and maps, whereas Emod was obtained for the same map/model pairs after applying coordinate bias or map sharpening/blurring.

### 3.6. A register shift in an RNA-dependent RNA polymerase complex

The RNA-dependent RNA polymerase (nsp12), is essential for replication of the SARS-CoV-2 viral genome. In a recent study, a cryo-EM structure of nsp12 with two accessory proteins nsp7 and nsp8 was determined at resolution 2.5 Å (pdb/emdb-id 7bv2/30210) providing valuable structural details on a mechanism of action of an antiviral drug remdesivir (Yin *et al*., 2020). Due to its importance, the structure has been carefully analysed by the structural biology community.

The original structure was shown to contain a number of potential problems that were quickly corrected in the updated PDB deposition (Croll, Williams *et al*., 2021). One of the problems was a register-shift of an isolated, and relatively short (15 amino-acids) α-helical fragment at the nsp12 C-terminus (Fig. 7a). A close inspection of the fragment shows that for example Threonine 912 in the original model is too small for the corresponding side-chain density, which can be better explained with Tyrosine 921 after shifting model register by 9 residues. This could not have been detected by a density-fit measure as the issue hasn’t resulted in prominent density outliers. Similarly, as the whole fragment is register-shifted there are no backbone geometry issues at the flanks of the shifted stretch. Nevertheless, the issue can be easily spotted with *checkMySequence* as a clear sequence-assignment outlier (Fig. 7a, red bar in the lower panel). Moreover, a new sequence assignment suggested in method’s text output could have been used to correct the issue, as it agrees with the updated coordinates (Fig. 8).

**Figure 7.**
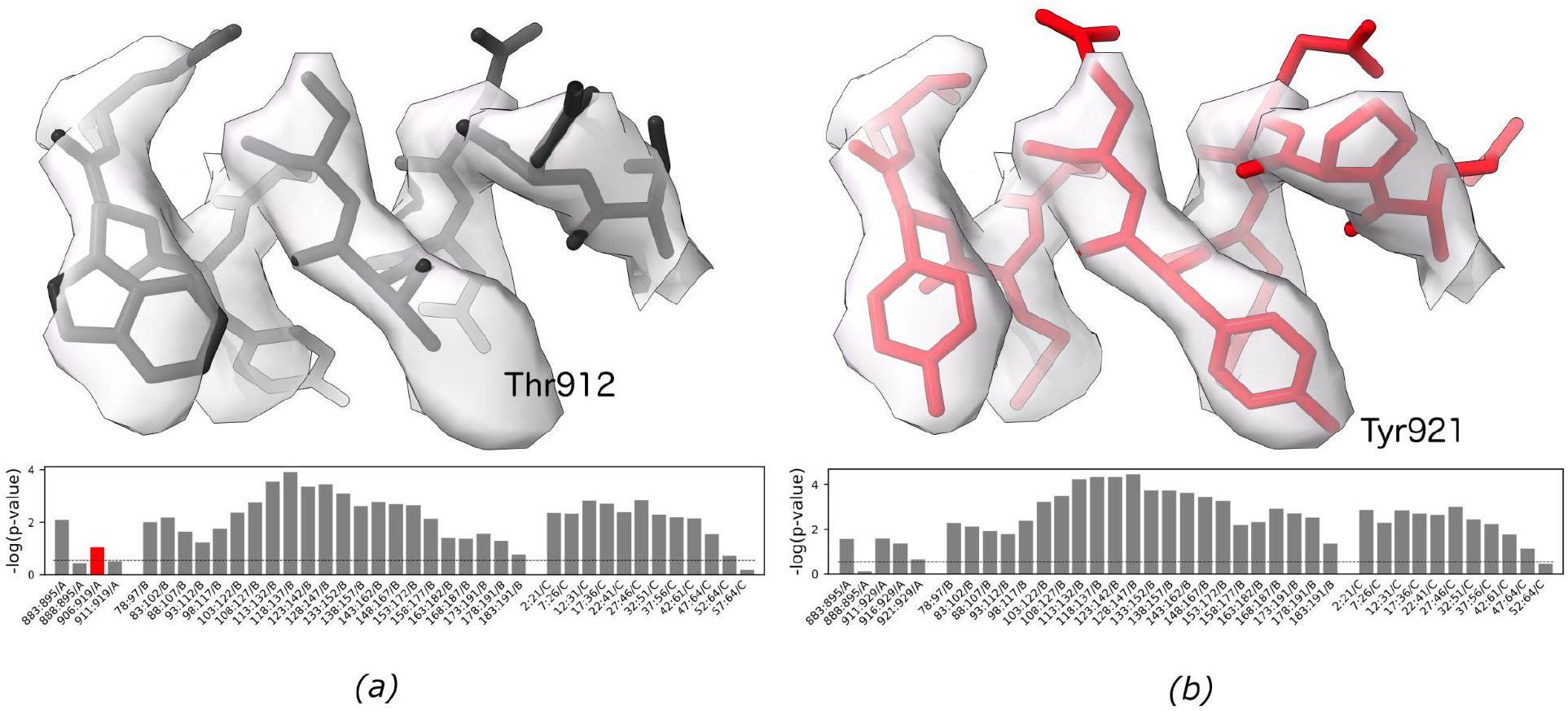
Fragment of an RNA-dependent RNA polymerase model (a) originally deposited version (pdb/emdb-id 7bv2/30210, residues A/908-917) and (b) model deposited by the authors after applying a register-shift error that was also suggested by *checkMySequence* (residues A/917-926). Bottom panels depict standard *checkMySequence* graphical output. Grey bars represent fragments for which the assigned sequence matches the model (dashed line). The red bar in panel (a) represents a fragment with register error part of which is shown in black. The bar-plots show −log(p-value) for the test-fragments; higher bars correspond to lower p-values and more reliable sequence assignments (be they correct or erroneous).

**Figure 8.**
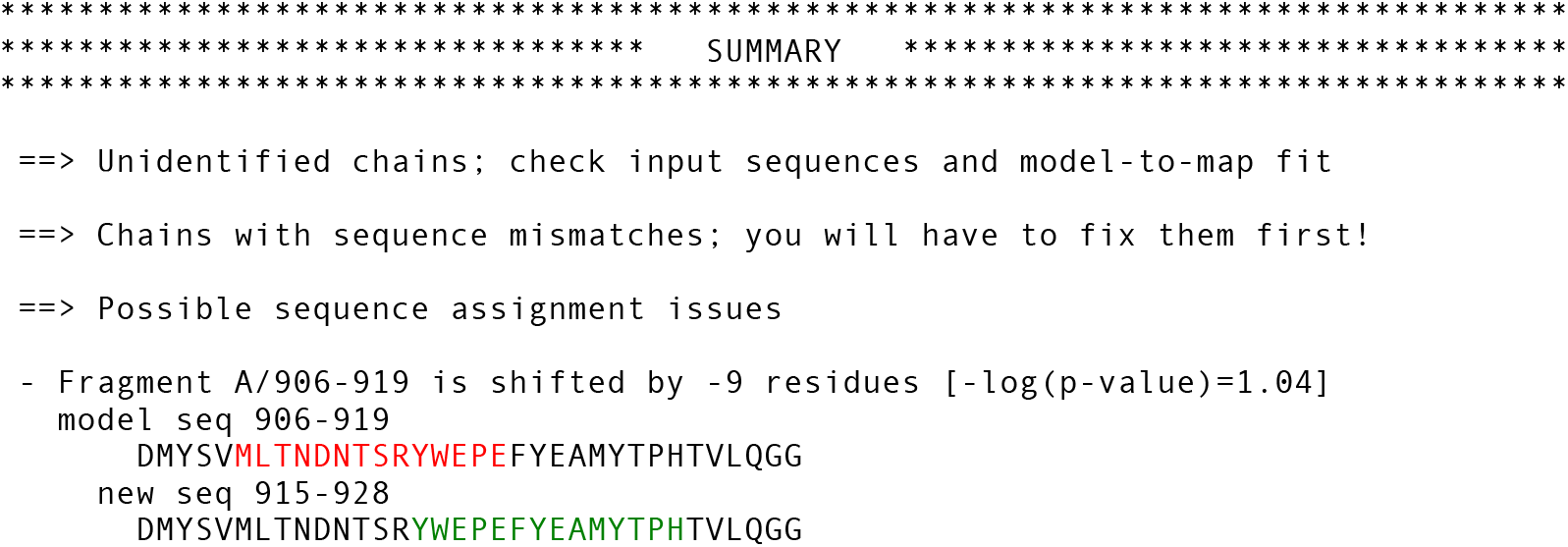
Standard text output of the *checkMySequence* for originally deposited version of RNA-dependent RNA polymerase model (pdb/emdb-id 7bv2/30210). The program prints possible issues with unidentified reference sequences, sequence mismatches, and register errors.

### 3.7. Cytoplasmic domain of a Transient receptor potential channel

In the benchmark set, we identified a number of clear outliers, where fragments were assigned sequences different from the reference model with low p-values. Particularly interesting was a cryo-EM structure of transient receptor potential cation channel subfamily C member 6 (TRPC6) determined at 3.8 Å resolution (pdb/emdb-id: 6cv9/7637, (Azumaya *et al*., 2018)). A number of fragments from this model form a clear, very low p-value cluster with assigned sequences different than reference (orange circles on Fig. 3b)

Despite the relatively low resolution, due to the lack of any close homologous structure, the original model was built *de novo* into the map using COOT, a very challenging task. The overall fold of the structure was later confirmed by a number of closely related, human TRPC6 structures determined at better resolutions. Nevertheless, we noticed a possible register-shift in a linker helical domain of the structure (Fig. 9a). As rebuilding the structure *de novo* would be very difficult owing to relatively low local-resolution, we decided to use a corresponding AlphaFold2 predicted structures database (Varadi *et al*., 2021) to interpret the map (id Q61143). Relevant region of the predicted model was fitted into the EM reconstruction using “Jiggle-Fit this molecule with Fourier Filter” tool and refined in real space with all-molecule self-restraints at 5 Å distance cut-off using COOT version 0.9.2-pre (Casañal *et al*., 2020) and additionally refined using phenix.real_space_refine version 1.18.2 (Afonine *et al*., 2018) with default parameters. Visual inspection of the new model showed a few clear signs of improvement in the match to the map (for example, the left-side end of α-helix presented on Fig. 9). Nevertheless, we noticed no prominent signs of sequence register improvement, for example for aromatic residues. The *checkMySequence* scores, however, improved significantly (Fig. 9, lower panels). Moreover, the sequence register of the new model confirmed the *checkMySequence* suggestions. We also noted that the new model sequence register agrees with a closely related (94% sequence identity) human structure that has been determined recently at 2.8 Å with much clearer side-chain densities in this region (pdb-id 6uz8, (Bai *et al*., 2020)). These results show that *checkMySequence* can be used for the interpretation of maps and model validation at resolutions where visual map interpretation is very challenging.

**Figure 8.**
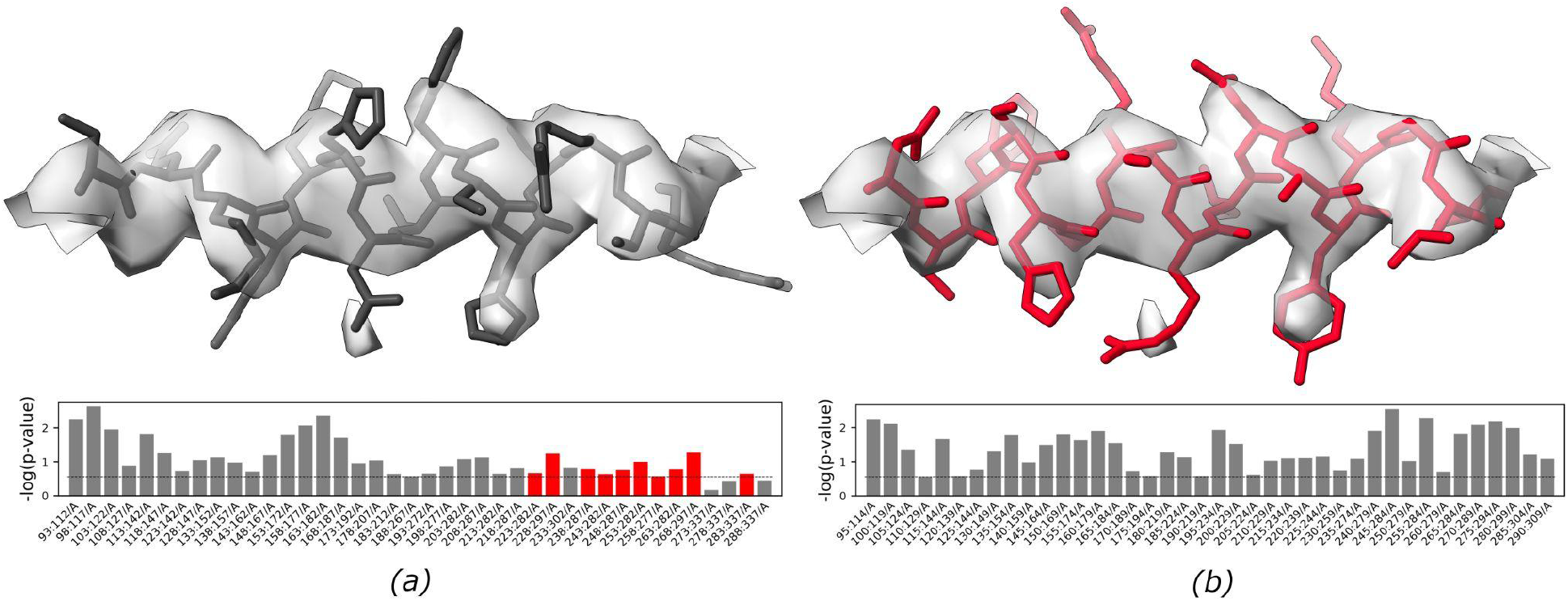
Fragment of mTRPC6 cytoplasmic domain at 3.8 Å resolution (a) deposited in PDB (pdb/emdb-id 6cv9/7637, residues A/258-272) and the same fragment rebuilt using AlphaFold2 prediction (Q61143, residues A/264-280, pLDDT of this region exceeds 90). Bottom panels depict corresponding *checkMySequence* graphical outputs. Grey bars represent fragments for which the assigned sequence matches the model or has a p-value higher than threshold (dashed line). Red bars on panel (a) represent fragments with plausible register errors, including the α-helix shown in black. The bar-plots show −log(p-value) for the test-fragments; higher bars correspond to lower p-values and more reliable sequence assignments.

## 4. Materials and Methods

### 4.1. Protein model benchmark set

Atomic model coordinates of macromolecular structures determined using cryo-EM were downloaded from PDB together with corresponding reconstructions and reference sequences as of 20 August 2021. Only structures determined at resolution 4 Å or better, with molecular weight below 500kDa, and half-maps available for download in EMDB (Velankar *et al*., 2016) were considered. In total, 796 structures fulfilled these criteria. For each of the half-map pairs local resolution maps were calculated using Resmap version 1.1.4 (Kucukelbir *et al*., 2014) with default parameters.

### 4.2. Protein chain test-fragment selection

Continuous protein-chain test-fragments were selected by shifting a “focus window” of fixed length in steps of 5 residues along all protein chains in a model. Only chains with at least 95% standard amino-acid residue content were considered. If multiple conformations of a residue were present in a model, only the first one was processed. The mean local resolution of the selected chain fragments was calculated for grid points of a corresponding local resolution map within 2 Å of any atom in a fragment. Random coordinate shift was applied for benchmarks only, to all model atoms independently ignoring any stereochemical restraints.

### 4.3. Map preprocessing

Before use, input map resolution was truncated in reciprocal space to 2.5 Å, to account for the absence of cryo-EM reconstructions determined at ultra-high resolution in the residue-type classifier training set implemented in *findMySequence*. Map blurring and sharpening was performed in reciprocal space and implemented in *checkMySequence* code using tools from CCTBX library.

### 4.4. Implementation and availability

The sequence validation program checkMySequence was developed based on routines implemented in *findMySequence*. It was developed in Python 3 with an extensive use of pytorch (Paszke *et al*., 2019), numpy (Oliphant, 2006), scipy (Virtanen *et al*., 2020), CCTBX (Grosse-Kunstleve *et al*., 2002) and CCP4 (Winn *et al*., 2011) libraries and utility programs. For making sequence database queries we use HMMER suite version 3.3.2. The program source code and installation instructions are freely available at https://gitlab.com/gchojnowski/checkmysequence under BSD-3 license.

## 5. Conclusions

Sporadic errors are an intrinsic part of the macromolecular model building process. Modeling in cryo-EM reconstructions seems to be particularly affected as target structures are usually large, map resolutions are heterogeneous, and the pressure to release models quickly is strong. Although a number of model validation tools have been developed to date, many of them specifically for cryo-EM map interpretation, register-shifts remain one of the most difficult problems to identify and correct.

Here we presented a new method, *checkMySequence* for the fully automated identification of register-shift errors in protein models built into cryo-EM reconstructions. The approach relies on a systematic assignment of short protein model test-fragments to the target sequence and works as a statistical test. If there is enough statistical evidence, the input-model sequence can be challenged, by providing a sequence that explains features of a map better. Although the strength of the approach in principle depends on factors like local map resolution, model quality, and a level of map sharpening or blurring, we have shown that these factors can be successfully compensated by an automated adjustment of a test-fragment length. This avoids the problem of bias related to user-provided parameters like map resolution. The method also removes the need for the user to understand and interpret validation scores: the results clearly state whether input model sequence assignment can be improved or not.

A limitation of this approach is its use of relatively long test-fragments, which are required to compensate for the low information content of cryo-EM maps. At lower local map resolutions, test-fragment length can be increased from default from 20 up to 60 amino acids. It means that a source of a sequence-register problem needs to be identified by a user within this range using interactive methods. It must be stressed however, that if a plausible register-shift is detected the method also suggests a new sequence assignment, which simplifies the error correction. The model errors can be conveniently rebuilt using tools available in a related method *findMySequence*. The requirement for long test-fragments, however, may also result in the method missing short, local register-shift errors, where a tracing issue is promptly compensated (e.g. a missing residue by an additional residue). Although the minimum length of a detectable register-shift stretch depends on resolution, it should be generally longer than 10 amino acids. Nevertheless, we show that the approach provides useful results at resolutions where a visual, residue-by-residue validation would be very challenging.

Apart from register-shift errors the program also checks the input model for sequence mismatches (single-residue differences between model and target sequence) and problems with residue numbering (e.g. continuous residue numbering ignoring gaps in a model) (Fig. 8). Another advantage of the presented method is its performance that makes it readily applicable to the analysis of very large models. For example, validation of a complete 70S ribosome structure at 3.0 Å resolution (pdb/emdb-id: 5we4/8814), with 58 protein chains and over 6,400 protein residues, takes less than three minutes on a basic laptop. As we have shown previously (Chojnowski, Simpkin *et al*., 2022), this structure contains several register-shift errors of the kind which could, in future in other cases, be avoided by use of *checkMySequence*. This is particularly important as detailed and exhaustive residue-by-residue analysis of such large models is rarely possible. In this context, the transparent graphical visualization (lower panels on Fig. 7 and 8) and command-line output of the validation results by *checkMySequence* should be of particular interest for users.

## Acknowledgements

We would like to thank all the microscopists who decided to deposit half-maps to EMDB. Although not strictly required, it was of great help while developing *checkMySequence*. Furthermore, we thank Daniel Rigden, and Alice Bochel for critical reading of the manuscript and very helpful comments. We also thank Filomeno Sanchez Rodriguez for valuable discussions and help with software testing.

